# ASPL-driven subunit exchange remodels VCP/p97 hexamers and is impaired by a multisystem proteinopathy mutation

**DOI:** 10.64898/2026.05.07.723621

**Authors:** Jingxuan Tang, Laxmikanta Khamari, Benjamin Dodd, Guoming Gao, Liuhan Dai, Stephanie L. Moon, Nils G. Walter

## Abstract

Valosin-containing protein (VCP/p97) is an essential homohexameric AAA+ ATPase that powers ubiquitin-dependent protein quality control by extraction and unfolding of clients for proteasomal degradation. Heterozygous, autosomal-dominant VCP missense mutants are associated with multisystem proteinopathy (MSP) through unclear molecular mechanisms. We developed a single-molecule pull-down assay to quantify VCP hexamer assembly and subunit exchange dynamics directly in human cell lysate. We show the common MSP-associated VCP variant R155H co-assembles with wild-type subunits to form heterohexamers. Wild-type VCP complexes readily undergo subunit exchange in cell lysates, but this exchange is markedly reduced for purified complexes in buffer. We identify the VCP-interactor ASPL as a selective mediator of monomer exchange that efficiently remodels wild-type, but only modestly exchanges R155H variants within multimers. Single-molecule kinetics analyses reveal ∼2-fold faster ASPL association with, and ∼4-fold slower dissociation from, wild-type VCP than R155H-VCP. We propose that ASPL-driven monomer exchange remodels VCP molecular machines to sustain proteostasis. The failure of ASPL-driven exchange of MSP variant monomers would be predicted to stabilize mutant VCP in assemblies, revealing a potentially targetable defect.

## Introduction

Neurodegenerative diseases such as amyotrophic lateral sclerosis (ALS) and frontotemporal dementia (FTD) present some of the most challenging medical problems in an aging population^1,2^. A distinctive feature of these disorders is the accumulation of misfolded proteins, underscoring the essential role of protein quality control in maintaining neuronal health^3,4^. The AAA+ ATPase valosin-containing protein (VCP, or p97) is a highly conserved, abundant hexameric protein that performs diverse cellular functions by extracting ubiquitinated substrates for recycling or degradation from cellular structures, including the endoplasmic reticulum (ER), stress granules, mitochondrial membrane, and ribosomes^5–14^. Through its unfolding activity, VCP regulates the stability, conformation, and subcellular localization of proteins.

Mutations in VCP are associated with multisystem proteinopathy (MSP), which encompasses a spectrum of degenerative conditions of the nervous, muscular, and skeletal systems. These degenerative conditions can overlap and include ALS, FTD, inclusion-body myopathy, and Paget’s disease of the bone^15–17^. MSP-associated VCP mutations are linked with disruptions in proteasomal degradation, stress granule clearance, autophagy, lysosomal homeostasis, and mitochondrial dynamics^13,14,18–27^. Despite the recognized clinical significance of missense VCP alleles, the precise molecular mechanisms by which VCP mutations trigger these pathologies are not fully understood, and no curative therapies are available.

In MSP associated with VCP mutations, affected individuals are typically heterozygous for the pathogenic VCP allele^15^. Reduced VCP function was recently linked to neurodegenerative phenotypes including tauopathy^28^. However, most MSP-associated alleles appear to act through toxic gain-of-function and/or dominant-negative effects, rather than simple haploinsufficiency. Pathogenic VCP mutations associated with MSP are inherited in an autosomal dominant manner^29^, and many studies report little to no change in steady-state VCP protein abundance^30–32^, although altered expression has been observed in some contexts^33,34^. Among MSP-associated VCP variants, R155H is most common^16,35^. Purified R155H-VCP homomers exhibit increased ATPase activity and enhanced substrate translocation *in vitro*^36–38^, supporting a hyperactive, potentially toxic state. Mutant and wild-type (WT) VCP subunits can co-assemble in bacterial expression systems when expressed from one plasmid^36^, and MSP mutations bias the VCP N-domain towards an “up” conformation^36,38^ that is associated with altered ATPase regulation. Because VCP hexamers are reported to be long-lived and resistant to subunit exchange in buffer^36^, a single mutant subunit incorporated into a mixed hexamer could, in principle, “poison” the complex. Since VCP unfolds substrates via a hand-over-hand mechanism, incorporation of a mutant subunit into the VCP hexamer is additionally expected to disrupt coordinated ATP-driven translocation and thereby affect substrate unfolding^39^. However, the extent, stoichiometry, and dynamics of WT–mutant VCP co-assembly in mammalian cells remain poorly defined.

VCP executes diverse proteostasis functions through ∼40 cofactors, many of which engage the N-terminal domain^17^. The N-terminal domain undergoes ATP/ADP-dependent conformational switching^40–42^ and MSP-associated mutations shift its conformational equilibrium, reshaping cofactor binding and downstream pathway engagement^43^. For example, the Npl4/Ufd1 heterodimer binds more tightly to mutant VCP, potentially rewiring ER-associated degradation, whereas UBXD1 binding is reduced, with possible consequences for organelle turnover pathways^36,44–47^. The VCP interactor ASPL (alveolar soft part sarcoma locus) is unusual among cofactors in that it can dissociate VCP hexamers into monomers and has been implicated in VCP post-translational regulation, including trimethylation and phosphorylation^48–50^. These observations raise the possibility that VCP assemblies are not static but can be actively remodeled via regulated subunit exchange within cells. Under this model, if a dominant-negative MSP-associated VCP allele is expressed at similar levels as WT VCP, ASPL would be predicted to mediate unbiased exchange between WT and mutant subunits and result in a ∼50:50 mixture of VCP within hexamers. However, immunoprecipitation/western blot analyses have suggested reduced ASPL association with R155H-VCP compared with WT VCP^50^. Whether this altered association impacts ASPL-mediated VCP disassembly or remodeling, and consequently VCP function, remains unknown.

Here, we establish single-molecule pulldown and photobleaching assays to quantify VCP hexameric assembly composition and dynamics directly in human (HeLa) cell lysate, and perform single-molecule kinetic measurements to resolve cofactor engagement. Using this platform, we examine co-assembly of WT and MSP mutant VCP into heterohexamers, quantify their stoichiometries, and test whether VCP undergoes subunit exchange. We further interrogate ASPL as a candidate remodeling factor and define how the most common MSP-associated VCP mutant allele alters ASPL binding and remodeling efficiency. Together, these experiments support a model in which ASPL-driven monomer exchange “refreshes” WT VCP machines—enabling cofactor reprogramming and limiting the persistence of dysfunctional subunits to maintain protein homeostasis—whereas impaired remodeling of mutant-containing assemblies provides a plausible, dominant route to proteostasis failure in MSP.

## Results

### Development of a single-molecule imaging assay for VCP hexamer assembly

Given the hexameric nature of VCP, it is plausible that WT and mutant subunits intermix during initial oligomer assembly, thereby modulating the structural and functional properties of the resulting complexes (Fig. 1a). We first established a single-molecule imaging assay for VCP hexamer assemblies by individually purifying recombinant Halo-tagged WT-VCP (WT_H_) and R155H mutant-VCP (MT_155-H_) from *E. coli* (Fig. 1b). These purified VCPs were subsequently labeled with HaloTag Alexa Fluor 660 and immobilized for single-molecule total internal reflection fluorescence (TIRF) imaging. We assessed via stepwise photobleaching assay whether the observed VCP assemblies indeed represented hexamers (Fig. 1c). Consistently, both WT and mutant VCPs exhibited an average of four discrete photobleaching steps (Fig. 1d). Accounting for the inherent labeling efficiency of ∼68%, these results indicate predominant hexamer formation. This observation suggests that, in isolation, both WT and mutant VCP proteins retain their ability to assemble into intact homohexamers, with the MT_155_ VCP exerting minimal impact on structural integrity. To further interrogate functional integrity, ATPase activity was measured: while WT VCP displayed typical enzymatic activity, the mutant demonstrated elevated ATPase activity, in alignment with previous studies^36–38^ (Fig. 1e). Collectively, these data demonstrate that VCP hexamer assembly can be quantified using single-molecule imaging assays and confirm that VCP hexamerization occurs despite the presence of the R155H mutation.

**Fig. 1.**
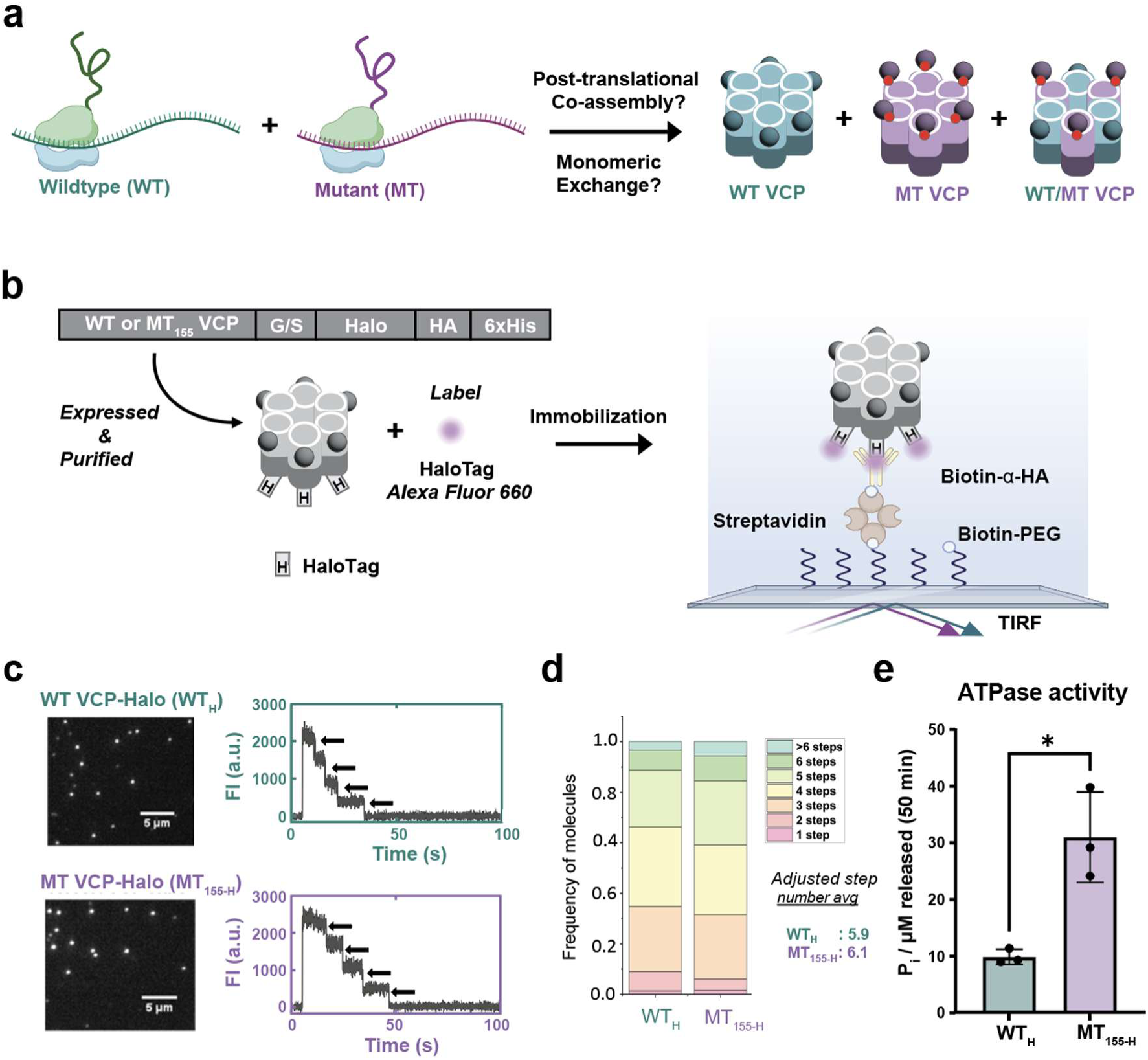
Wild-type and R155H mutant VCP form hexamers with comparable stoichiometry. **a** Schematic showing the heterogeneous expression systems in which WT and mutant monomers are co-expressed and potentially co-assemble in oligomer proteins. **b** Experimental schematic for single-molecule analysis: WT VCP–Halo (“WT_H_”) or R155H VCP–Halo (“MT_155-H_”) was expressed in *E. coli*, purified, surface-immobilized, and imaged by single-molecule total internal reflection fluorescence (TIRF) microscopy. **c** Representative frames from single molecule experiment described in (a): fluorescence images showing HaloTag–Alexa Fluor 660–labeled WT-VCP–Halo (top) and R155H VCP–Halo (bottom); corresponding fluorescence intensity trajectories (right) illustrate stepwise photobleaching used to infer monomer stoichiometry per hexamer. **d** Distribution of monomer counts per VCP hexamer derived from photobleaching step analysis. **e** ATPase activities of purified WT-VCP–Halo and R155H-VCP–Halo measured under the indicated conditions. The average +/- SD from *n* = 3 independent experimental replicates is shown. The statistical significance of differences was determined using Student’s *t*-test ( \**p* < 0.05).

### Wild-type and mutant VCP co-assemble upon co-expression

Given the hexameric nature of VCP, it is known that WT and mutant subunits intermix during bacterial co-expression^36^, thereby modulating the structural and functional properties of the resulting complexes (Fig. 1a). To test whether this observation translates into the human system, we co-expressed SNAP-tagged WT-VCP (WT_S_) and Halo- tagged R155H-VCP (MT_155-H_) in HeLa cell lysate, employing fluorophores with minimal spectral overlap, SNAP-Surface 549 and HaloTag Alexa Fluor 660, to differentially label the two monomers (Fig. 2a). We leveraged TIRF microscopy to directly visualize and quantify the formation of VCP hexamers. To do so, we incubated biotin-PEGylated coverslips with streptavidin, followed by biotinylated anti-HA antibody to pull down the fluorescently labeled HA tagged VCPs (all VCP constructs were C-terminally fused to an HA tag for immobilization) expressed from plasmids in coupled transcription/translation reactions in the lysate.

**Fig. 2.**
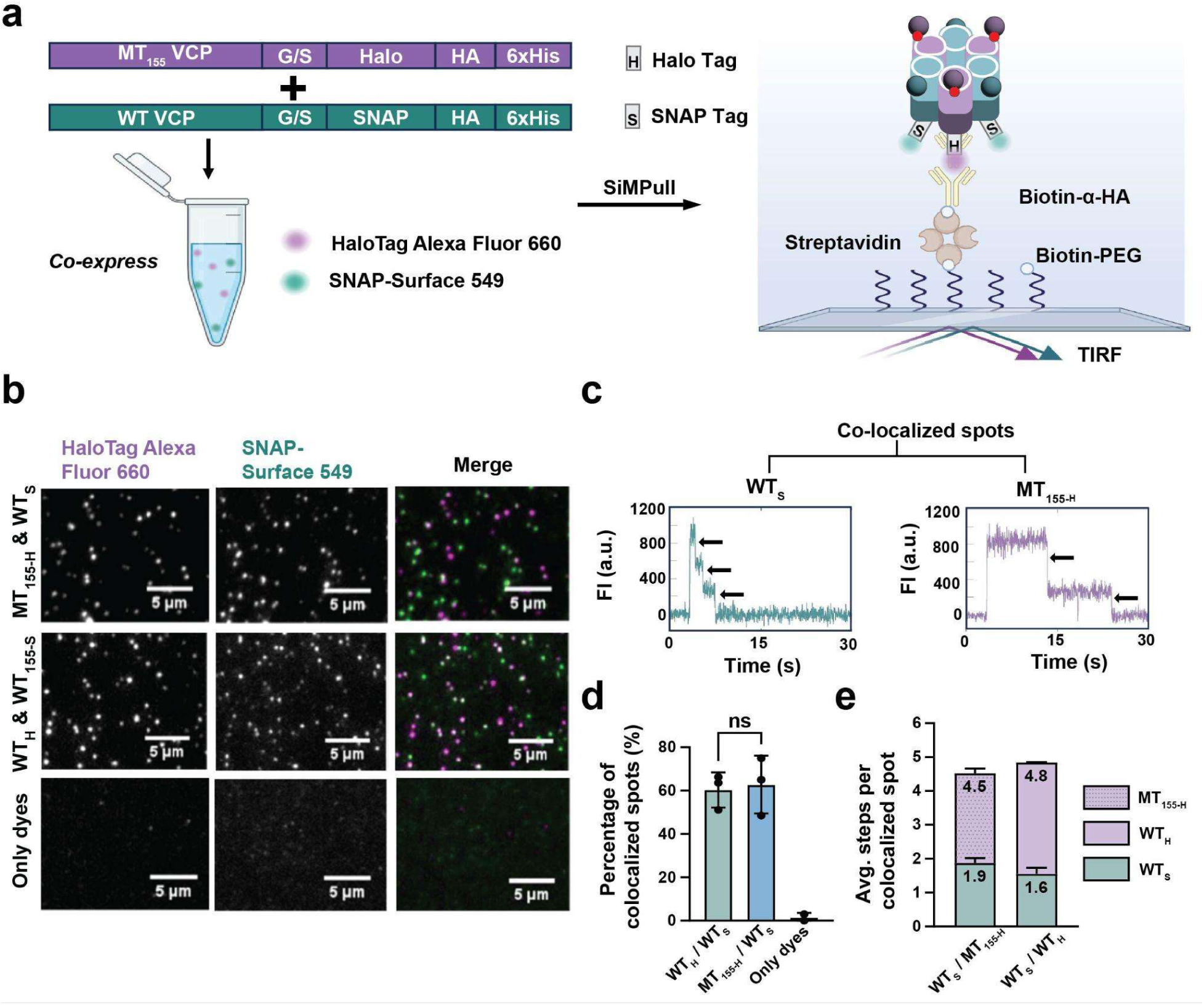
Co-assembly of WT and R155H VCP monomers into mixed hexamers in Hela lysate. **a** Schematic of the co-expression experiment. WT-VCP–SNAP and R155H-VCP–Halo plasmids were co-transfected into HeLa lysate, labeled with SNAP-Surface 549 and HaloTag–Alexa Fluor 660, respectively, and imaged by single-molecule TIRF microscopy. **c** Representative TIRF images showing R155H VCP–Halo (“MT_155-H_”; Alexa Fluor 660; left), WT VCP–SNAP (“WT_S_”; SNAP-Surface 549; middle), and the merged channels (right). **d** Representative intensity–time trajectories from colocalized Halo- and SNAP-labeled spots showing discrete photobleaching steps. **e** Colocalization frequencies for WT/WT and WT/R155H co-expression conditions. **f** Subunit stoichiometry within colocalized complexes, expressed as the ratio of Halo-channel to SNAP-channel photobleaching steps for WT/WT and WT/R155H samples. The mean +/- SD from *n* = 3 independent experimental replicates is reported. The statistical significance of differences was determined using the Student’s *t*-test.

We observed discrete foci of WT_S_ and MT_155-H_ representing one or more VCP monomers adhering to the coverslip (Fig. 2b). A majority (∼60%) of VCP particles were two-color heterohexamers containing both WT and mutants (Fig. 2d). Control lysates containing free dyes alone produced negligible background (Fig. 2d), underscoring that the observed colocalization of WT and mutant VCP monomers reflects VCP monomer co-assembly. These findings reveal that mutant VCP co-assembles with WT VCP into oligomers as previously suggested by bulk gel filtration of bacterial expression assays^36^.

Having established that WT and MT_155_ VCP subunits are capable of co-assembly, our next objective was to assess whether the presence of a mutation within a VCP monomer alters assembly efficiency by evaluating the resultant stoichiometry of subunit incorporation within oligomeric complexes. By co-expressing equimolar amounts of WT_S_ and either WT-Halo (WT_H_) or MT_155-H_ plasmids in HeLa lysate, we could distinguish and enumerate subunit incorporation based on the TIRF-monitored photobleaching of the distinct fluorescent labels (Fig. 2b,c). In the WT_S_/MT_155-H_ experiments, the average photobleaching steps detected were approximately 2 for WT and 2.5 for the mutant subunits (Fig. 2e). As a control, the WT_S_/WT_H_ co-expression condition exhibited averages of 1.5 and 3 steps, respectively (Fig. 2e). The combined average step count across both conditions was around 4.5; less than the expected six steps for fully labeled hexamers, which can be attributed to the inherently sub-stoichiometric (∼68%) labeling efficiency observed in purified protein controls. Notably, we found that WT and MT_155_ VCP monomers co-assembled to a similar degree into heterohexamers (Fig. 2e), indicating that the R155H mutation does not impair assembly into heterohexamers.

### ASPL facilities subunit exchange of VCP hexamers

Because WT and mutant VCP monomers can co-assemble, we next asked whether co-assembly occurs during the initial formation of the hexameric complex, or through monomer exchange between homohexameric complexes. This distinction is important, as the high stability of the hexameric complexes in buffer^36^ could suggest the potential for mutated monomers to poison pre-formed WT assemblies for long periods of time. We sought to investigate whether this rigidity persists in the more complex environment of the human cell lysate. We first assessed whether mixing independently purified WT and mutant VCP proteins in HeLa lysate resulted in mixed WT-mutant VCP co-assemblies (Fig. 3a). TIRF images were acquired (Fig. 3b), and colocalization analysis revealed that WT and mutant VCP co-assemble at approximately 30% (Fig. 3c). In contrast, we did not observe co-assembly of WT and mutant VCP when we mixed each recombinantly purified protein in a buffer (Fig. 3c). These results suggest that an activity in mammalian lysate promotes dynamic VCP subunit exchange.

**Fig. 3.**
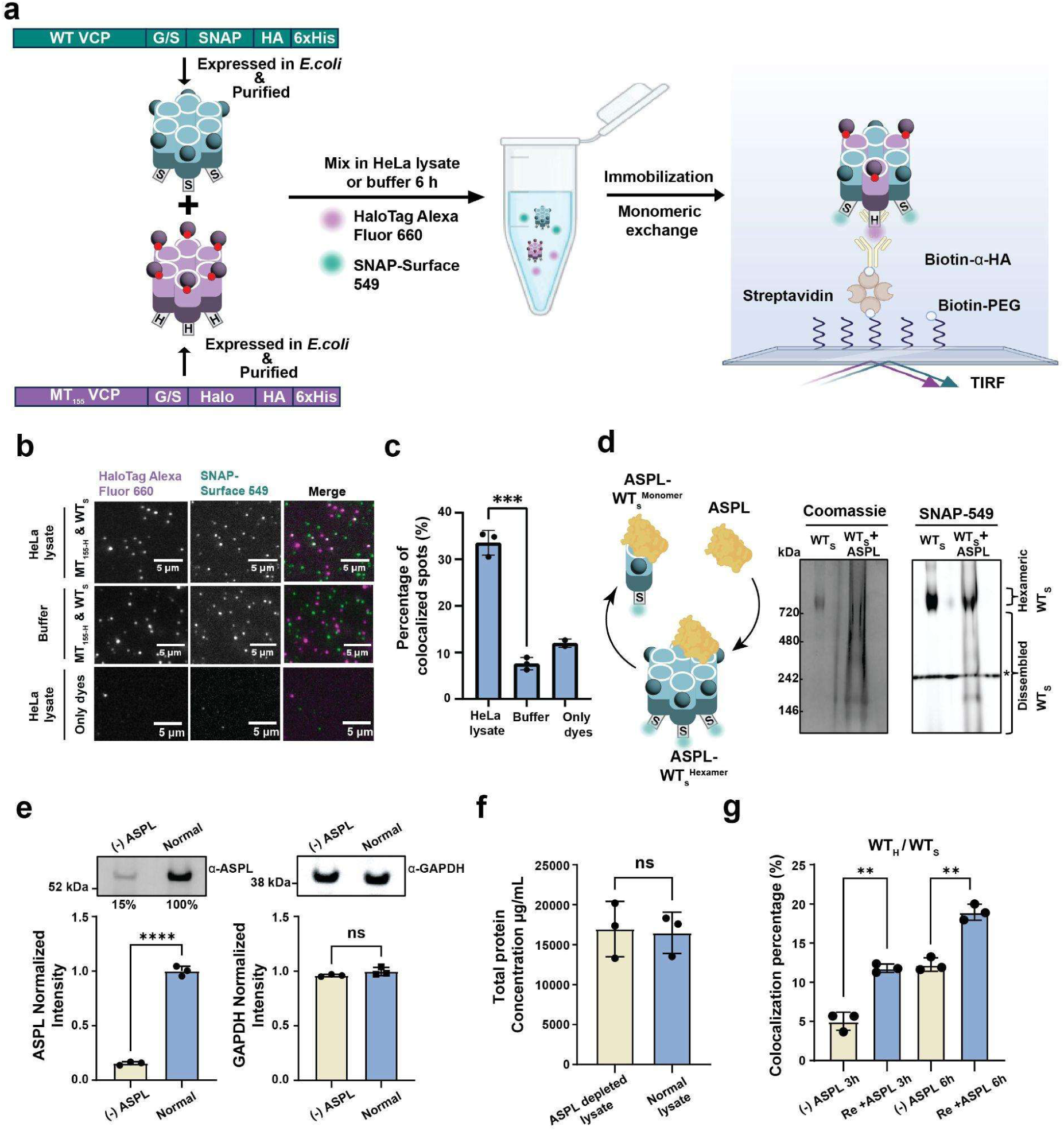
Recombinant WT and R155H VCP complexes exchange subunits in HeLa lysate. **a** Schematics of experiment testing whether WT VCP and R155H VCP exchange monomers. WT-VCP-SNAP (“WT_S_”) and R155H-VCP–Halo (“MT_155-H_”) were expressed separately in *E. coli* and purified, incubated in HeLa lysate for 6 hours, then fluorescently labelled with Alexa Fluor 660 (Halo) and SNAP-Surface 549 (SNAP). Complexes were immobilized on coverslips and imaged by single-molecule TIRF microscopy. Co-localization of the two fluorophores was used as a readout of subunit exchange between WT and R155H assemblies. **b** Representative single-molecule images of labelled R155H VCP–Halo (left; magenta) and WT VCP–SNAP (middle; green); merge shown at right. **c** Quantification of colocalized WT and R155H spots in HeLa lysate and in buffer, with free-dye control. **d** Blue native PAGE of recombinant VCP and ASPL (left, Coomassie stain; right, fluorescence). * indicates the bands present in all lanes arising from components of the commercial native gel buffer. **e** Immunoblot (top) and quantification (bottom) showing ∼85% ASPL depletion by immunoprecipitation; GAPDH serves as a loading control. **f** Total protein concentration in HeLa lysate before and after ASPL immunodepletion. **g** Co-localization between WT-VCP-SNAP and WT-VCP-Halo in ASPL-depleted and ASPL-repleted HeLa lysates after 3 h and 6 h incubation, as marked in the figure. The mean +/- SD from *n* = 3 independent experimental replicates is shown. The statistical significance of differences was determined using Student’s *t*-test (\*\*\*\**p* < 0.0001, \*\*\**p* < 0.001, \*\**p* < 0.01).

While many factors could contribute to monomer exchange, we focused on elucidating whether the VCP cofactor ASPL facilitates VCP monomer exchange, since ASPL can dissociate VCP hexamers^49^ (Fig. 3d). We first confirmed that ASPL can dissociate WT hexamers by mixing purified WT_S_ with ASPL in solution and analyzing the products via non-denaturing gel electrophoresis, revealing a loss of the hexameric species consistent with ASPL’s disassembly activity (Fig. 3d). We then tested whether ASPL mediates WT_H_/WT_S_ monomer exchange by first immuno-depleting ∼85% of endogenous ASPL from HeLa lysate (Fig 3e) and verifying that the lysate otherwise remained unperturbed, by analyzing GAPDH levels (Fig. 3e) and total protein concentration (Fig. 3f). Next, we performed single molecule colocalization analyses, which revealed that WT_H_/WT_S_ co-assembly was significantly diminished upon ASPL depletion, but could partially be restored by supplementation with recombinant ASPL (Fig. 3g).

Together, these results demonstrate that ASPL is necessary and sufficient for WT VCP subunit exchange between multimeric assemblies.

### Single-molecule kinetic analysis reveals ASPL differentially regulates wild-type and mutant VCP assemblies

Recent results of co-immunoprecipitation experiments from cell cultures indicate that ASPL interacts with R155H VCP with reduced affinity compared to WT VCP^50^. We therefore used single-molecule imaging to examine whether ASPL differentially affects WT and MT_155_ monomer exchange. We first further characterized the impact of either augmentation or depletion of ASPL from HeLa lysates on the colocalization of WT and mutant VCP monomers. Using western blotting to quantify the concentration of endogenous ASPL in HeLa lysate compared to purified ASPL of defined concentrations, we found 34 pg/μL of endogenous ASPL in our HeLa lysate (Supplementary Fig. 1). We then supplemented the lysate with either 34 pg/μL purified ASPL or BSA–the latter as a control for nonspecific effects–and assessed whether VCP co-assembly changed. Notably, only the colocalized fraction of WT_S_/WT_H_ assemblies increased upon ASPL addition relative to BSA, whereas the colocalized fractions of WT_S_/MT_155-H_ and MT_155-S_/MT_155-H_ were unchanged (Fig. 4a). These data suggest that ASPL preferentially mediates monomer exchange among WT VCP subunits, likely reflecting its stronger affinity for WT VCP relative to mutant forms. We quantified colocalization extent after mixing purified proteins in HeLa lysate for 10 minutes, observing that the colocalized fractions are ∼5% across the WT_S_/WT_H_, WT_S_/MT_155-H_, and MT_155-S_/MT_155-H_ pairings (Fig. 4a), similar to the results for buffer, indicating negligible monomer exchange during this initial window. We designate this sample as the effective zero time point.

**Fig. 4.**
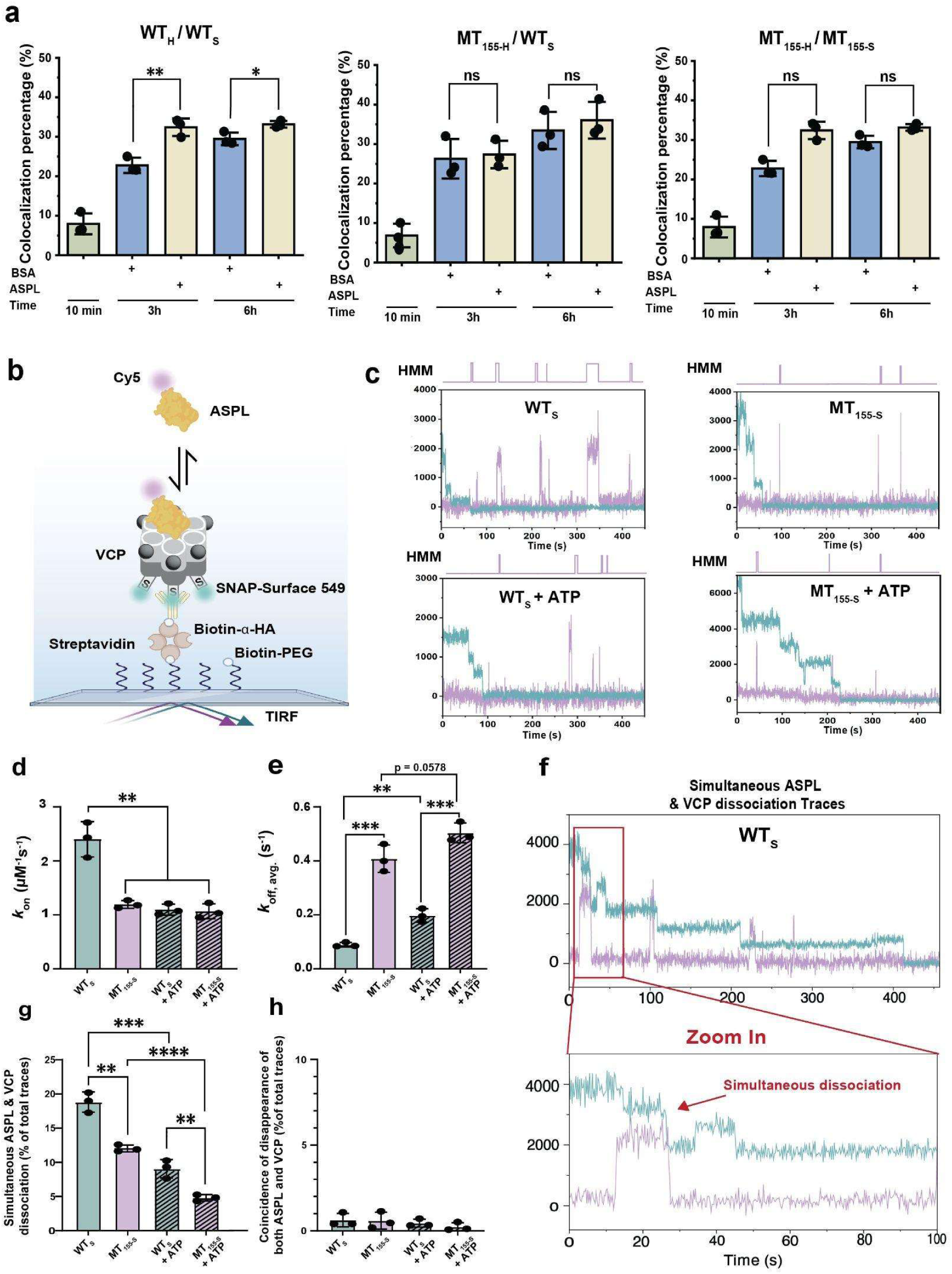
ASPL binds tighter with WT VCP than R155H VCP and modulates VCP subunit exchange. **a** Colocalization percentages of purified WT and R155H (MT_155_) VCP. WT_S_/WT_H_, WT_S_/MT_155-H_, and MT_155-S_/MT_155-H_ were incubated in HeLa lysate for 10 min, 3 hours, and 6 hours with or without an additional excess amount of ASPL or BSA, as indicated. **b** Schematic of the single-molecule TIRF experiment with Cy5-labeled ASPL interacting with immobilized WT_S_ or MT_155-S_. **c** Representative kinetic traces of ASPL (purple), interacting with WT_S_/MT_155-S_ VCP (cyan) in the absence of ATP, and in the presence of ATP. The cyan traces are photobleaching steps of SNAP-tagged VCP which shows the oligomeric states of VCP. **d** The association rate constant, k_on_, and **e** dissociation rate constant, k_off_ , of ASPL interacting with WT_S_ or MT_155-S_ VCP with or without ATP, as indicated. **f** Trace in which the signals of ASPL and WT_S_ VCP monomer disappear in the same frame. **g** Percentage of traces in which the signals of ASPL and VCP monomer disappear in the same frame for WT or mutant VCP, with or without ATP. **h** Coincidence of disappearance of signal from both ASPL and VCP after photobleaching and dissociation. The mean +/- SD from *n* = 3 independent experimental replicates is reported. The statistical significance of differences was determined using Student’s *t*-test (\*\*\*\**p* < 0.0001, \*\*\**p* < 0.001, \*\**p* < 0.01, \**p* < 0.05).

To elucidate the mechanistic underpinnings of the differential ASPL-WT and MT_155_ interactions, we conducted single-molecule kinetic assays using surface-immobilized WT_S_ or MT_155-S_ and incubated with Cy5-labeled ASPL, in the presence or absence of ATP (Fig. 4b). Because VCP’s N-terminus shifts upward in the ATP-bound state, and downward in the ADP-bound state, adding hydrolyzable ATP is expected to reveal how nucleotide-dependent conformational changes influence VCP’s interaction with ASPL. Real-time fluorescence traces were acquired to characterize ASPL binding and dissociation from VCP, with VCP photobleaching step analysis used to filter for traces corresponding to VCP assemblies larger than three subunits (Fig. 4c). Using quantitative kinetic analysis of cumulative dwell time distributions in the unbound and bound states, we determined the association (*k*_on_; Fig. 4d) and dissociation (*k*_off_; Fig. 4e) rate constants, respectively, for ASPL with either WT or MT_155_ VCP (see also Supplementary Fig. 2). Notably, ASPL exhibited an almost two-fold higher binding rate constant with WT than MT_155_ (Fig. 4d), suggesting facilitated recognition of WT VCP.

Introduction of ATP markedly decreased *k*_on_ for WT, consistent with a model wherein ATP-induced upward movement of the N-terminal domains impedes ASPL access by partial occlusion of its binding site^36^. Consistent with this model, MT_155_ VCP–which intrinsically favors upward N-terminal positioning–did not exhibit this ATP-dependent reduction in *k*_on_, supporting the notion that its more prevalent upward conformation drives the mutant’s diminished ASPL interaction.

Further analyses revealed a nearly four-fold lower ASPL *k*_off_ for WT than MT_155_ (Fig. 4e). Adding ATP increased the *k*_off_ for WT more than MT_155_, consistent with the more prominent ATP-induced structural rearrangements of WT^36^ and suggesting steric competition between the upward N-terminal domain and ASPL. Combining the association and dissociation kinetics, we calculated equilibrium dissociation constants (*K_d_* = *k*_off_/*k*_on_) that suggest that ASPL exhibits a ∼9-fold higher affinity toward WT than MT_155_ VCP (Supplementary Fig. 2).

Our kinetics data suggest that ASPL binds to MT_155_ VCP with lower affinity than WT VCP, leading to diminished exchange of the former. To further test this idea, we asked whether we could observe ASPL-driven monomer removal in real time. To do so, we closely inspected our time traces (Fig. 4f) and quantified the fraction in which ASPL signals disappeared simultaneously with either WT or MT_155_ signal (Fig. 4g). Strikingly, ∼18.8 ± 1.2% of ASPL/WT traces exhibited at least one simultaneous signal disappearance of ASPL and VCP, whereas only ∼12.1 ± 0.4% of ASPL/MT_155_ traces did (Fig. 4g), far exceeding an expected coincidental co-disappearance of ASPL and VCP of <1% (Fig. 4h). As expected from the increased N-terminal upward conformation, addition of ATP significantly decreased the fraction of traces in which signals disappear simultaneously for WT and MT_155_ to 8.9 ± 1.1% and 4.8 ± 0.4%, respectively. Given the persistent difference in the proportion of simultaneous signal disappearance between ASPL/WT and ASPL/MT_155_, our results suggest that ASPL facilitates WT monomer exchange more frequently than that of MT_155_. As a consequence, MT_155_ is predicted to persist in ASPL-containing heterohexamers (Fig. 5), consistent with the observed reduced ASPL-driven monomer exchange.

**Fig. 5.**
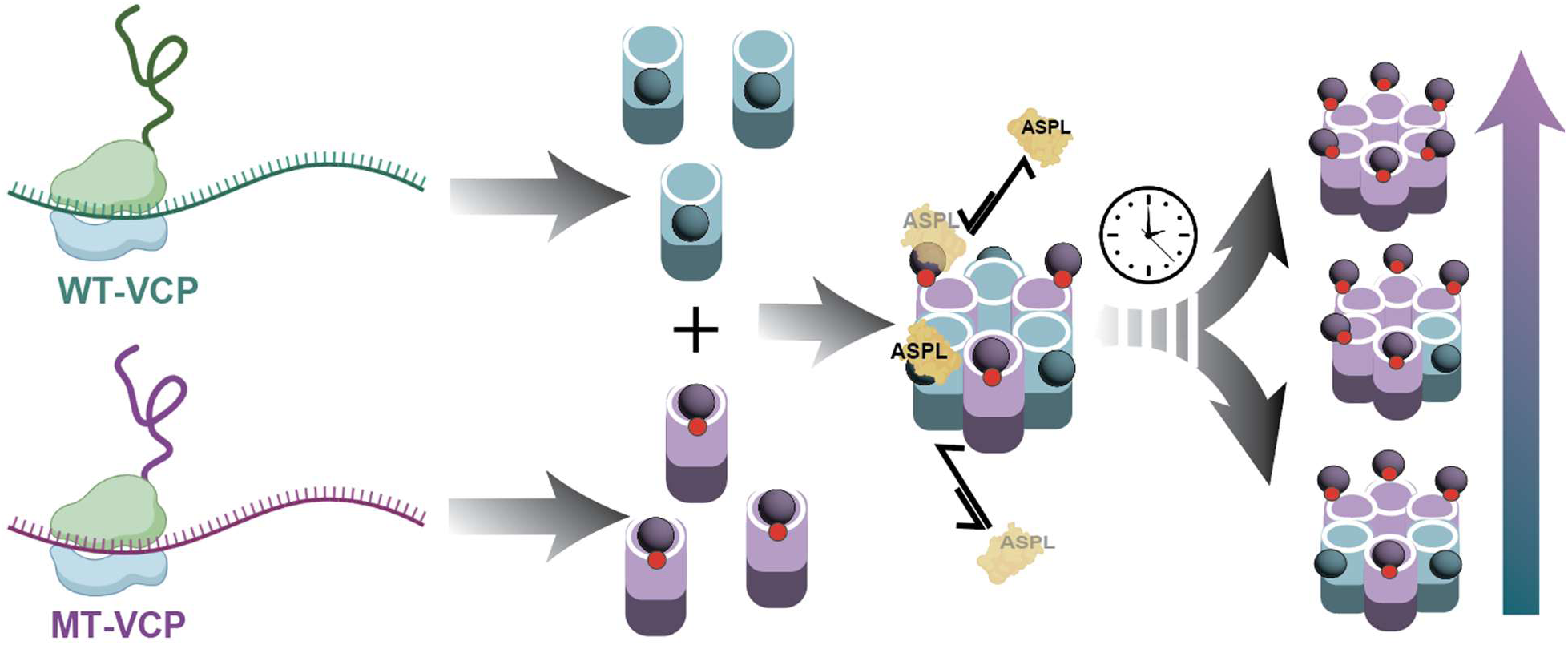
Model of ASPL as an accelerator of monomer exchange in wild-type VCP, with limited binding and exchange of the MSP mutant R155H-VCP leading to the accumulation of mutant components in assembled machines over time.

## Discussion

Our study establishes single-molecule assays that reveal how WT and MSP-associated mutant subunits co-assemble into human VCP and how ASPL dynamically regulates hexamer composition under near-cellular conditions (Fig. 5). Using a single-molecule pull-down strategy, we immobilized HA-tagged VCP from HeLa lysate on PEGylated coverslips and directly visualized hexamer formation (Fig. 1) and subunit exchange (Fig. 2). These experiments identify ASPL as a key exchange factor that promotes remodeling of WT hexamers (Fig. 3) but refreshes mutant-containing assemblies less efficiently (Fig. 4). Beyond defining an exchange mechanism for VCP, our approach provides a general framework for interrogating multimeric machines at the level of individual assemblies and offers new insight into MSP pathogenesis.

Our data indicate that VCP hexamers in human lysate behave as dynamic assemblies that exchange subunits, in sharp contrast to purified VCP in buffer, which shows little exchange (Fig. 3c). This supports a model in which cellular factors actively renew VCP complexes and that ASPL is a principal driver of this remodeling. In heterozygous MSP, such renewal creates a route to dominant-negative effects: ASPL-mediated opening and reassembly enable mutant monomers to enter pre-existing WT hexamers, generating heterohexamers with impaired performance. The preferential remodeling of WT assemblies further implies that renewal is biased, leaving mutant-containing complexes comparatively refractory to refreshment (Fig. 5).

This exchange-based model also reconciles the apparent stability of VCP hexamers inferred from their *in vitro* behavior^36–38,40–42^. If hexamers were effectively irreversible once assembled, mixed WT/mutant complexes in heterozygous cells would be expected to arise predominantly at initial assembly, likely near sites of translation. Under that scenario, transcription timing and local concentration effects would favor mostly WT or mostly mutant hexamers rather than extensively intermixed assemblies. Instead, we observe ready WT–mutant co-assembly in lysate, and independently purified hexamers become mixed within lysate, indicating that post-assembly remodeling actively interconverts VCP complexes in a cellular milieu.

We therefore propose a hexamer renewal model for VCP homeostasis in which cells maintain proteostasis not only by regulating VCP abundance and cofactor usage, but also by periodically resetting VCP assemblies through ASPL-driven disassembly and subunit exchange. Such renewal could help remove cumulative damage or modifications that accrue with stress, aging, and disease^51–53^; as well as facilitate the incorporation of programmed post-translational modifications into monomers of active VCP complexes^30,48,50,54–67^. Notably, ASPL need not selectively recognize a damaged subunit; instead, repeated opening and reassembly would refresh assembly state and cofactor occupancy without requiring wholesale turnover of the complex.

A key implication of our findings is that ASPL refreshes WT hexamers more efficiently than mutant-containing hexamers. If ASPL-mediated disassembly is part of a normal renewal cycle, then WT monomers would enter that cycle readily, whereas mutant monomer components would be longer-lived and remodeled less frequently. In heterozygous cells, this asymmetry could engender dominant-negative effects, even if WT is predominantly expressed (Fig. 5): WT monomers would be liberated more often, increasing opportunities for mutant monomers to incorporate during reassembly and sustain heterohexamer formation.

This framework connects naturally to MSP as an age-linked degenerative disease^29,68–70^. If VCP assemblies progressively accumulate damage or become trapped in maladaptive functional states, renewal would become increasingly important over time. MSP mutations such as R155H would compromise renewal by weakening ASPL engagement, slowing association and accelerating dissociation, thereby reducing disassembly and exchange and biasing the system toward persistence of mutant-containing hexamers. The resulting imbalance could progressively burden VCP-dependent protein quality control, consistent with adult-onset degeneration^29,68–70^. Our discovery that ASPL more stably remodels WT than mutant hexamers explains emerging observations that VCP disease mutants disrupt ASPL-mediated VCP phosphorylation and G3BP1 extraction, resulting in persistent stress granules that could contribute to ALS/FTD pathogenesis^50^.

Therapeutically, our results suggest strategies aimed at restoring renewal capacity and limiting exchange-driven “poisoning.” One approach would be to enhance ASPL-mediated remodeling—for example, by stabilizing ASPL engagement with mutant-containing assemblies or by shifting VCP conformational equilibria toward states permissive for productive ASPL action. A complementary approach would be to reduce mutant incorporation during exchange, including allele-selective reduction of mutant VCP or interventions that bias reassembly toward WT-enriched complexes. More broadly, our findings imply that MSP can be viewed as a failure of regulated VCP machine renewal, a defect that may be therapeutically tractable.

These considerations also bear on efforts to treat MSP by modulating VCP ATPase activity. Incorporation of mutant subunits into a hexamer is expected to disrupt intersubunit coordination during ATP-driven remodeling, compromising substrate processing and downstream pathways. ATPase inhibitors (for example NMS-873, ML240, and CB-5083) have shown benefits in some models^31,67^, but MSP is typically heterozygous so that WT and mutant VCP co-exist and assemble together. Nonselective inhibition may therefore suppress residual WT activity within heterohexamers, potentially exacerbating loss of function and contributing to cytotoxicity. Our observation of extensive mixed-oligomer formation underscores the need for interventions that preserve or restore WT VCP activity while counteracting mutant effects.

Finally, our data strengthen the view that cofactor interactions are coupled to VCP assembly dynamics, with implications for MSP. MSP mutations alter N-domain orientation and remodel cofactor affinities^40–42^—enhancing binding to some partners (e.g., UFD1–NPL4, VCF1) while weakening that to others (e.g., UBXD1 and ASPL)^36,44–46^. How monomer exchange reshapes cofactor occupancy, and how many WT subunits are required within a hexamer to sustain specific cofactor-driven functions, remain important open questions. Although we focused here on R155H, the assay framework should be readily extended to additional disease alleles (including A232E, R155C, R159H, R191Q, and L198W)^10^. More generally, the principles uncovered here are likely applicable to other oligomeric protein quality-control machines implicated in neurodegeneration. Our observation of compositional dynamics in VCP adds to a growing recognition that many canonical ‘machines’ in the cell are not compositionally fixed, but instead undergo continual subunit and partner turnover as they function^71^. This raises the possibility that similar exchange-driven remodeling will be found across other AAA+ ATPases and oligomeric quality-control assemblies, where activity and specificity emerge from competing kinetic exchange processes rather than static occupancy and conformational changes alone. If so, measuring exchange rates and compositional lifetimes may become as important as structural snapshots for defining mechanisms and for understanding how disease mutations rewire function. As a paradigm, future work integrating additional cofactors and post-translational modifications—including ASPL-mediated trimethylation and phosphorylation—should further clarify how renewal efficiency of VCP/p97 is tuned and how its failure contributes to disease^48,50^.

## Methods

### Plasmid generation and protein purification

Human WT or mutant VCP cDNA was cloned into a bacterial expression vector encoding a C-terminal Halo-tag, HA-tag, and a six-histidine tag, separated by a flexible G/S linker (GGGGSGGGGSGGGGSGGGGSGGGGSG). Plasmids were transformed into BL21(DE3) *E. coli* (New England Biolabs C2527H). Starter cultures were inoculated overnight in 5 mL LB with 0.25 mg Kanamycin, then used to seed 1 L LB with 50 mg Kanamycin at OD_600 = 0.01. Cultures were grown at 37°C, 250 rpm, to OD_600 = 0.6, induced with 0.5 mM IPTG (Millipore Sigma I6758), and incubated for 14 h at 25°C. Cells were harvested by centrifugation (4°C, 5000g, 30 min), lysed in buffer (50 mM Tris-HCl pH 8, 300 mM NaCl, 0.5 mM β-mercaptoethanol, protease inhibitor; Millipore Sigma 4693132001), and clarified by centrifugation at 20,000g. The supernatant was filtered (0.45 μm), mixed with 20 mM imidazole, and passed over a Ni-NTA column (Qiagen 30210). Bound protein was eluted with sequential imidazole steps (50–300 mM), and peak fractions were pooled, buffer-exchanged into Buffer A (20 mM HEPES pH 7.4, 250 mM KCl, 1 mM MgCl_2_), concentrated with Amicon 10k filter (Millipore Sigma UFC801024), flash-frozen in liquid nitrogen, and stored at –80°C with 5% glycerol and 0.5 mM TCEP (Sigma Aldrich C4706). The size and purity of the purified protein was verified using denaturing PAGE.

### *In vitro* VCP transcription/translation assays

For expression of VCP in human lysates, the *in vitro* transcription/translation reaction contained 10 μL HeLa cell extract, 2 μL accessory proteins, 4 μL reaction mix (Thermo Scientific 88881), 500 ng each of WT-SNAP and WT-Halo, or 500 ng each of WT-SNAP and mutant-Halo VCP plasmids, 3.66 μM of HaloTag Alexa Fluor 660 Ligand (Promega G8471) and/or SNAP-Surface 549 (New England Biolabs S9112S) dyes, then water added to 20.5 μL total volume. Incubation was for 6 h at 30°C prior to imaging, unless otherwise indicated.

### Single-Molecule Fluorescence Microscopy

Single-molecule fluorescence imaging was performed using an Oxford Nanoimager (ONI) benchtop platform operating in objective-based total internal reflection fluorescence (TIRF) mode. The system was outfitted with dual-band emission filters and a Hamamatsu ORCA-Flash4 V3 sCMOS camera (instrument specifications: https://oni.bio/nanoimager/). All experiments utilized a 100× oil immersion objective with a numerical aperture of 1.4, and autofocus was maintained via the system’s integrated Z-lock module. Experiments were conducted at a controlled temperature of approximately 25 °C, regulated by the ONI’s built-in temperature controller. Prior to imaging, the microscope was equilibrated to the set temperature, and the water bath temperature was adjusted as needed after laser activation to maintain a stable environment. TIRF illumination was achieved at an incident angle of ∼53.5°. SNAP-Surface 549 and HaloTag Alexa Fluor 660 Ligand or Cy5 dye-labeled proteins were excited at 532 nm and 640 nm.

### Single-molecule colocalization and photobleaching assays

Glass coverslips were coated with PEG/biotin-PEG (100:1), blocked with 5 mg/mL BSA (Blocker BSA Thermofisher 37525 diluted with buffer A) for 5 min, and functionalized with 1 mg/mL streptavidin (Thermo Fisher S888), followed by 0.01 mg/mL biotinylated anti-HA antibody incubation (Invitrogen 26183-BTIN) at 4 °C for 10min. Co-expressed and labeled VCP in lysate was diluted in buffer A with 1 mg/mL BSA and incubated for 40 min. The coverslip was washed with buffer A with 1 mg/mL BSA, and imaged under an oxygen-scavenging buffer (PCD 125 mM, PCA 12.5 mM, Trolox 8.33 mM). All washing steps are 3 times post incubation with buffer A and 1mg/mL BSA. Imaging was performed on an Oxford Nanoimager at 30 frames per second, for a total of 4,000 frames per channel. A high-power laser (640nm and 532nm) was used to photobleach the fluorescent molecules.

Spot localization was carried out using the TrackMate plugin in ImageJ, and subsequent analysis for dual-color colocalization was performed using custom Python scripts (distance threshold: 200 nm; calibration: 117 nm/pixel). Colocalization was quantified as the fraction of total molecules exhibiting both green and red fluorescence within each field of view. Photobleaching traces were extracted from the colocalized spots, and individual photobleaching steps were counted manually due to the high signal-to-noise ratio of the traces.

### Labeling efficiency estimation of individually purified VCP

The 6 µM individually purified WT-Halo VCP was labeled with 24 µM HaloTag Alexa Fluor 660 Ligand for 1 hour at 25 °C. The labeled proteins were denatured with NuPAGE™ LDS Sample Buffer (4X) (Thermofisher NP0007) and 1 mM DTT, and run on a NuPAGE denaturing gel. An Amersham™ Typhoon™ biomolecular imager was used to scan for fluorescent intensity. Labeling efficiency was calculated by determining the ratio of dye associated with the protein to free dye and normalized to total dye concentration.The estimated labeling efficiency was 68%.

### Monomer exchange assay

30 nM each of purified VCP WT-SNAP and mutant VCP-HaloTag proteins were co-incubated in HeLa lysate (10 μL HeLa cell extract) with 2 μL accessory proteins and 4 μL reaction mix (Thermo Scientific 88881) or Buffer A control, together with 3.66 μM each of HaloTag Alexa Fluor 660 ligand (Promega G8471) and SNAP-Surface 549 (New England Biolabs S9112S). Water was added to a final volume of 20.5 μL, and reactions were incubated for 6 h. Where indicated, exogenous ASPL (Abcam ab167886) or BSA (0.56 μg) was added to double the endogenous ASPL concentration. After incubation, VCP complexes were immobilized and analyzed as above for colocalization detection using the Oxford Nanoimager.

### ASPL-mediated VCP dissociation detection via non-denaturing gel

SNAP-Surface 549 labeled VCP WT-SNAP at 3.2 μM (monomer concentration, 0.53 μM as hexamer) was incubated with either 3.2 μM purified ASPL protein (Abcam ab167886) or buffer alone for 15 minutes. The samples were then loaded onto a Blue Native PAGE gel (Thermo Fisher BN1002BOX) and imaged using an Amersham™ Typhoon™ biomolecular imager.

### ASPL depletion and western blotting

For western blotting: 1.28 μL of Biotinylated anti-ASPL antibody (0.78 mg/mL) (Novus Biologicals NBP1-28709B) or 1.28 μL PBS buffer was incubated with 10 μL Dynabeads™ Streptavidin Magnetic Beads (Thermo Fisher 65601) for 30 min at room temperature. The beads were separated from solution and washed with PBS 3 times. 6.4 μL HeLa lysate mix (4 μL HeLa lysate + 0.8 μL accessory protein + 1.6 μL reaction mix) (Thermo Scientific 88881) was then added for ASPL depletion (or no ASPL antibody control). Samples were incubated overnight at 4 °C. Beads were removed and total protein content measured in lysate before and after depletion by the BCA assay (Thermo Fisher 23225). Western blotting for ASPL and GAPDH was done with ASPL/TUG Monoclonal Antibody (Thermo Fisher MA5-44891) and anti-GAPDH (Proteintech 60004-1-Ig) on denaturing PAGE gels, using secondary antibodies IRDye 800CW Donkey anti-Mouse IgG Secondary Antibody (LI-COR 926-32212) and IRDye 680RD Donkey anti-rabbit IgG Secondary Antibody (LI-COR 926-68073).

For the monomer-exchange ASPL depletion assay, the same protocol was performed at a 2.5× proportionally larger volume. Biotinylated anti-ASPL antibody (3.2 μL; 0.78 mg/mL; Novus Biologicals NBP1-28709B) or 3.2uL PBS buffer was incubated with Dynabeads™ Streptavidin Magnetic Beads (25 μL; Thermo Fisher 65601). After washing, 10 μL HeLa lysate, 2 μL accessory proteins, and 4 μL reaction mix (Thermo Scientific 88881) was added. Purified VCP proteins were then added to lysates as described in the monomer-exchange assay.

### Fluorescent labeling of ASPL

ASPL was non-specifically labeled with Cy5-NHS ester dye (Lumiprobe, Cat. #13020) at a 1:10 molar ratio of protein to dye in a total volume of 50 µL labeling buffer (1X PBS, pH 7.4; Thermo Fisher Scientific, Cat. #10-010-049), yielding a final protein concentration of approximately 10 µM. The reaction was incubated at room temperature for 1 h. Unreacted dye was removed by two sequential separations using Zeba™ Spin Desalting Columns with a 7 kDa molecular weight cutoff (Thermo Fisher Scientific, Cat. #89882). Labeling efficiency was quantified spectrophotometrically using a NanoDrop instrument by measuring absorbance at ∼280 nm (protein) and ∼650 nm (Cy5 dye). The labeling efficiency was approximately 100%, corresponding to a 1:1 molar ratio of Cy5 dye to ASPL protein. The labeled ASPL was supplemented with 50% glycerol, flash-frozen, and stored at −80 °C.

### Preparation of slide surfaces for single-molecule microscopy

Glass coverslips (No. 1.5, 24 × 50 mm²; VWR, Cat. #48393-241) were functionalized with a mixture of mPEG-SVA and biotin-PEG-SVA (10:1 ratio; Laysan Bio, Cat. #MPEG-SVA-5000-1g and #BIO-PEG-SVA-5K-100MG, respectively), following established protocols^72,73^. To preserve the PEGylation, the treated coverslips were wrapped in aluminum foil and stored in a nitrogen-filled chamber for up to four weeks. Prior to use, six sample chambers were assembled on each coverslip by cutting approximately 2 cm from the broader end of standard micropipette tips (Thermo Fisher, Cat. #02-682-261), discarding the narrow section, and placing the wide end directly onto the PEG-coated surface. The perimeters of the chambers were then sealed with epoxy adhesive (Ellsworth Adhesives, Cat. #4001).

### Single-molecule experiments for binding assay of ASPL with VCP

Biotin-PEG–passivated coverslips were first washed three times with 1× T-50 buffer. Streptavidin (1 mg/mL) was then introduced into each chamber and incubated for 10 minutes to facilitate biotin-streptavidin binding. The chambers were subsequently rinsed with a washing buffer consisting of 1× PBS (pH 7.4; Thermo Fisher Scientific, Cat. #10-010-049) containing 1 mg/mL BSA. Following this, ∼10 nM biotinylated anti-HA antibody was added and incubated for 20 minutes at room temperature, after which the chambers were washed again with washing buffer. Next, 20 pM of HA-tagged VCP was applied and incubated for an additional 20 minutes at room temperature, followed by another three washes using the same buffer.

For imaging, an imaging buffer was employed, consisting of 1× PBS (pH 7.4; Gibco), an oxygen scavenging system comprising 5 mM 3,4-dihydroxybenzoic acid (Fisher, #AC114891000), 0.05 mg/mL protocatechuate 3,4-dioxygenase (Sigma-Aldrich, #P8279-25UN), and 1 mM Trolox (Fisher, #218940050), and 10 nM Cy5-ASPL. After successful immobilization of VCP protein on the coverslip, the chambers were washed with a washing buffer, the imaging buffer was added, and single-molecule experiments were initiated immediately.

### Data acquisition and analysis for ASPL binding with VCP by colocalization assays

Data acquisition and analysis were performed using a custom MATLAB script. The intensity-versus-time traces were extracted from the raw datasets using this code. Traces were then selected manually based on the following criteria: at least four-step photobleaching events of SNAP-Surface 549, which is labeled with VCP, to confirm the presence of VCP homo-hexamers, as well as Cy5 fluorescence spikes exceeding ten times the background intensity. Traces exhibiting binding events were subsequently idealized with a two-state model (bound and unbound) employing the segmental k-means algorithm implemented in QuB. The dwell times corresponding to the bound (τ_on_) and unbound (τ_off_) states of ASPL were determined from these idealized traces. Cumulative distributions of these dwell times were then generated and fit with single or double exponential functions in Origin, yielding lifetime estimates for each state. The dissociation rate constant (k_off_) was calculated as the reciprocal of τ_on_, while the association rate constant (k_on_) was determined by dividing the reciprocal of τ_off_ as well as ASPL concentration used during data acquisition.

### Data Availability

All data supporting the findings of this study are available within the paper and its Supplementary Information.

## Acknowledgements

We thank Katelyn Green and Andreas Schmidt for their assistance. We appreciate funding support from Chan Zuckerberg Initiative Collaborative Pairs grants 2022-250629 to SLM and 2022-250725 to NGW, R35GM146711 to SLM, and a PPG Summer Research Fellowship and Rackham Graduate Student Research Grant to JT.

## Author contributions

Project conceptualization: BD, NGW, SLM; Study Design: JT, LK, NGW, SLM; Experimental conceptualization: BD, JT, LK, NGW, SLM; Data Curation: JT, LK; Formal analysis: GG, JT, LK, LD, NGW, SLM; Funding acquisition: NGW, SLM; Investigation: JT, LK; Methodology: JT, LK; Project administration: NGW, SLM; Resources: BD, JT, LK; Software: GG, LD; Supervision: NGW, SLM; Validation: JT, LK; Visualization: JT, LK; Writing - original draft: JT LK; Writing - review & editing: JT, LK, NGW, SLM.

## Competing interests

The authors declare no competing interests.

## Declaration of Generative AI use

Generative AI was used to edit grammar and improve writing clarity. No new content was generated, and all scientific interpretations and conclusions remain those of the authors.

## Notes

### Competing Interest Statement

The authors have declared no competing interest.

