## Supplementary Figures 1 and 2 for "ASPL-driven subunit exchange remodels VCP/p97 hexamers and is impaired by a multisystem proteinopathy mutation"

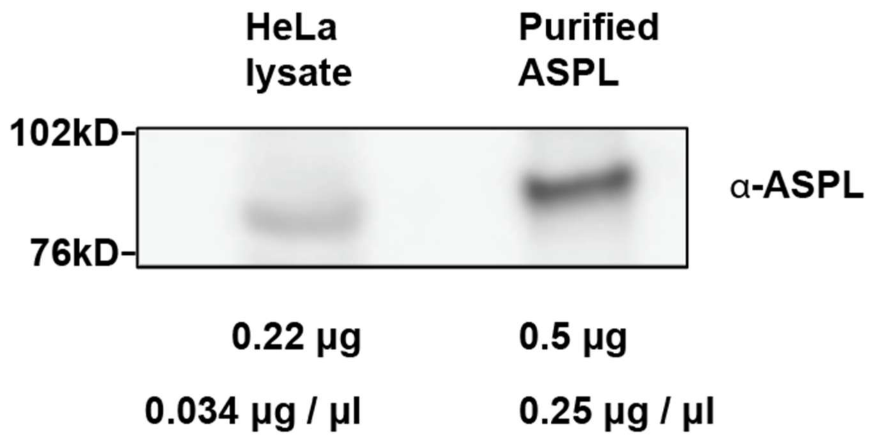


**Supplementary Fig. 1|** HeLa lysate ASPL concentration quantification by western blot with purified ASPL, as a standard.

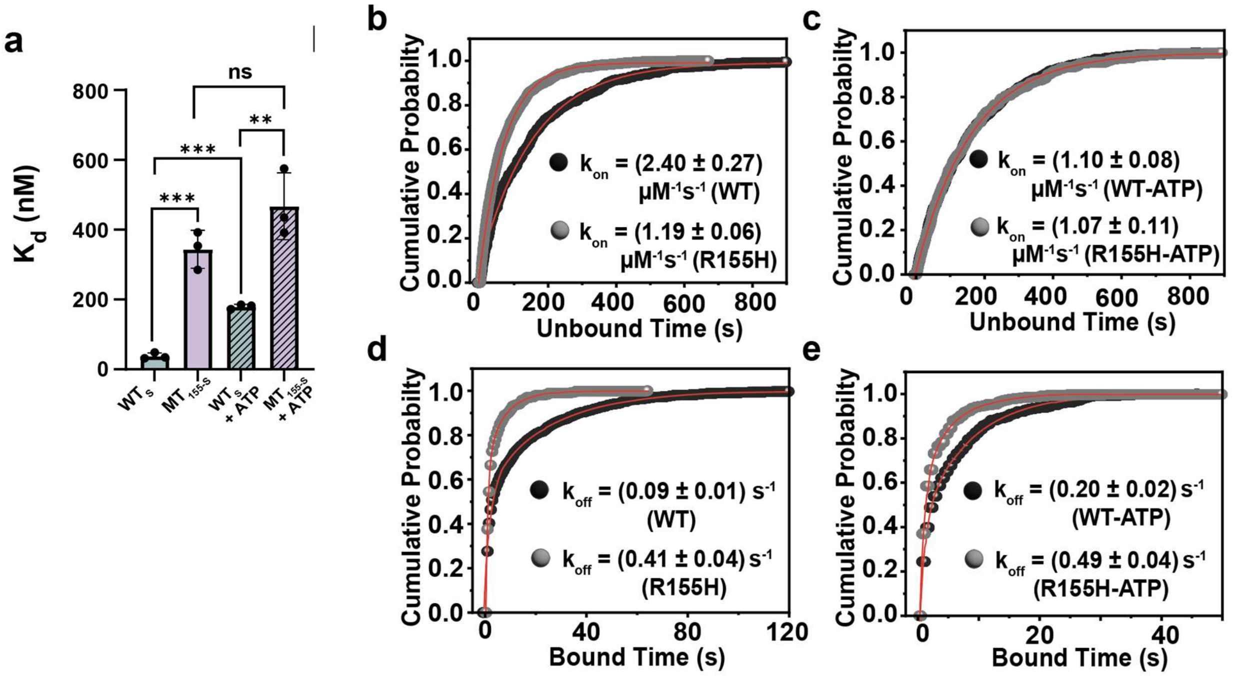


**Supplementary Fig. 2| Binding kinetics and equilibrium dissociation constant $K_d$, of ASPL to WT and R155H VCP. a** Equilibrium dissociation constant, Kd (estimating from $k_{on}/k_{off}$) of ASPL interacting to WT or R155H VCP with or without ATP, as marked in the figure. **b-c** Cumulative probability distributions of unbound dwell times for Cy5-labeled ASPL interacting with WT and R155H-VCP: **(a)** in the absence of ATP and **(b)** in the presence of ATP. **d-e** Cumulative probability distributions of bound dwell times for Cy5-labeled ASPL interacting with WT and R155H VCP: **(d)** in the absence of ATP and **(e)** in the presence of ATP. Error bars are the SD of the mean from *n* = 3 independent experimental replicates. The statistical significance of differences was determined using the Student's *t*-test (***$p < 0.001$, **$p < 0.01$, *$p < 0.05$).
